# ^13^C Electron Nuclear Double Resonance Spectroscopy Shows Acetyl-CoA Synthase Binds Two Substrate CO in Multiple Binding Modes and Reveals the Importance of a CO-Binding ‘Alcove’

**DOI:** 10.1101/2020.06.23.165407

**Authors:** Christopher D. James, Seth Wiley, Stephen W. Ragsdale, Brian M. Hoffman

## Abstract

EPR and Electron Nuclear Double Resonance spectroscopies here characterize CO binding to the active-site A cluster of wild-type (WT) Acetyl-CoA Synthase (ACS) and two variants, F229W and F229A. The A-cluster binds CO to a proximal Ni (Ni_p_) that bridges a [4Fe-4S] cluster and distal Ni_d_. An alcove seen in the ACS crystal-structure near the A-cluster, defined by hydrophobic residues including F229, forms a cage surrounding a Xe mimic of CO and is suggested to ‘cradle’ this CO. Previously, we only knew WT ACS bound a single CO in the A_red_-CO intermediate, here seen as forming Ni_p_(I)-CO with CO on-axis of the d_z_^2^ odd-electron orbital (g_⊥_>g_‖_∼2). The two-dimensional field-frequency pattern of 2K-35 GHz ^13^C-ENDOR spectra collected across the A_red_-CO EPR envelope now reveals a second CO bound in the d_z_^2^ orbital’s equatorial plane. This WT A-cluster conformer dominates the nearly-conservative F229W variant, but ^13^C-ENDOR reveals a minority “A” conformation with (g_‖_>g_⊥_∼2) characteristic of a ‘cloverleaf’ (eg. d_x_^2^-_y_^2^) odd-electron orbital, and with Ni_p_ binding two, apparently ‘in-plane’ CO. Disruption of the alcove through introduction of the smaller alanine residue in the F229A variant diminishes conversion to Ni(I) ∼tenfold and introduces extensive cluster flexibility. ^13^C-ENDOR shows the F229A cluster is mostly (60%) in the “A” conformation, but with ∼20% each of the WT conformer and an “O” state in which d_z_^2^ Ni_p_(I) (g_⊥_>g_‖_∼2) surprisingly lacks CO. This paper thus demonstrates the importance of an intact alcove in forming and stabilizing the Ni(I)-CO intermediate in the Wood-Ljungdahl pathway of anaerobic CO and CO_2_ fixation.

## Introduction

Here we provide new insights into the enzymatic control of carbon monoxide binding to acetyl-CoA synthase (ACS), the key enzyme in anaerobic carbon dioxide fixation by the Wood-Ljungdahl Pathway (**Eq 1**). The catalytic cycle of ACS is unique in involving a series of nickel-bound organometallic intermediates^1^ This sequence of steps resembles that of the rhodium-based Monsanto industrial acetic acid process.^2,3^

In the biochemical pathway (**Scheme 1**), CO is generated by CO dehydrogenase (CODH), which forms a tight complex with ACS and catalyzes the two-electron reduction of CO_2_ (**Eq 2**) at a Ni-[4Fe4S] cluster (the C-cluster). The C-cluster is linked to an electron transfer chain including two nearby [4Fe-4S] clusters (B-and D-clusters). The CO then migrates through a 75 Å hydrophobic tunnel from CODH to the active site of ACS, called the A-cluster, a dinickel-[4Fe4S] cluster in which a proximal Ni (Ni_p_) bridges the cluster and a distal Ni_d_.^4-7^

There has been controversy over whether the ACS catalytic mechanism proceeds through paramagnetic or diamagnetic intermediates.^1, 8-10^ The diamagnetic mechanism proposes a formally Ni_p_(0) species that undergoes carbonylation or methylation to form Ni_p_(0)-CO or Ni_p_(II)-CH_3_.^10^ In contrast, the paramagnetic mechanism proposes an active Ni_p_(I) species, which forms Ni_p_(I)-CO or Ni_p_(III)-CH_3_. In support of the paramagnetic mechanism, reduction and carbonylation of as-isolated CODH/ACS generates a kinetically-competent EPR-active species originally called the NiFeC complex, denoted A_red_-CO here, is detected.^11-14^ In supportive work on model systems, a nickel-substituted azurin model exhibits Ni(I)-CO, Ni(III)-CH_3_ and acetyl-Ni species.^15-17^ The A_red_-CO species, which is the focus of studies here, has been spectroscopically characterized by EPR,^14, 18^ ENDOR,^13^ Mössbauer,^19^ IR,^20^ and EXAFS spectroscopic methods.^21^

According to the paramagnetic mechanism (**Scheme 1**), when Ni_p_ binds CO, it forms a Ni_p_(I)-CO intermediate in which the unpaired electron is highly delocalized among the components of the A-cluster, including the bound CO.^12-14, 21^ During turnover, Ni_p_ also binds Coenzyme A (CoA) and a methyl group from a corrinoid iron-sulfur protein (CFeSP), then catalyzes C-C (forming acetyl-Ni) and C-S bond synthesis to generate the key metabolic precursor, acetyl-CoA (**Eq. 3**).

The existence of a tunnel connecting the active sites of CODH and ACS was first revealed by kinetic measurements,^4, 22^ in one case by a demonstration that labeled ^14^CO generated from ^14^CO_2_ does not exchange with unlabeled ^12^CO in solution as it undergoes conversion by ACS to [1-^14^C]-acetyl-CoA.^4^ Subsequent crystallographic studies using Xe as a stand-in for CO visualized gas molecules all along the interprotein tunnel, caged in specific sites by large hydrophobic residues. The most compelling gas-binding site is an ‘alcove’ surrounded by hydrophobic residues, seen with a Xe 3.9 Å above the A-cluster proximal Ni_p_, which bridges the distal Ni_d_ and the [4Fe-4S] center (See **Fig 1**). ^5^ Photolysis of the Ni(I)-CO dissociates CO; the rebinding activation energy is remarkably low (∼1 kJ/mol), suggesting the photodissociated CO remains trapped within the alcove, which facilitates its rebinding to Ni_p_.^23^

**Figure 1.**
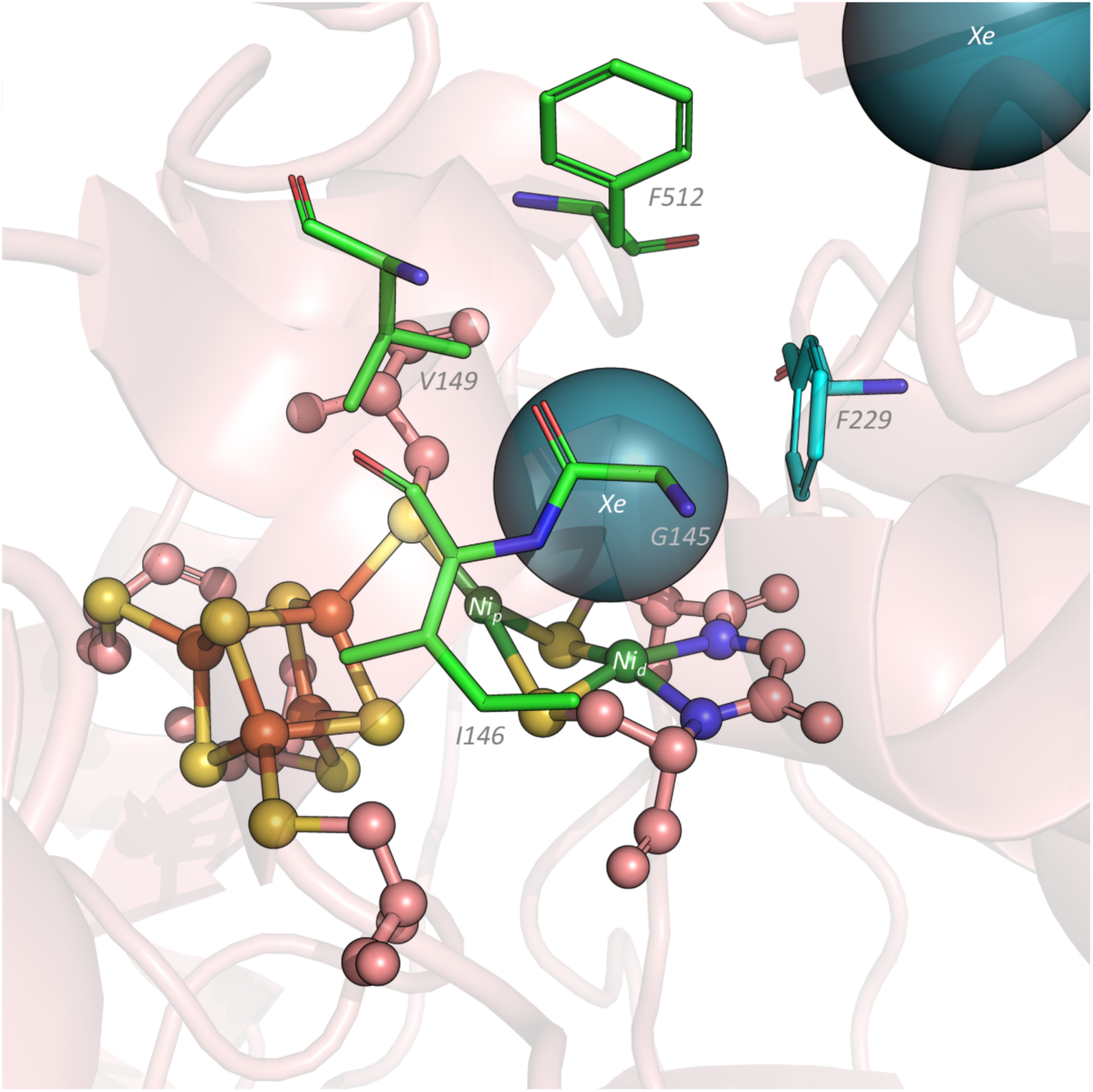
X-ray structure of the ACS A-cluster under high pressures of xenon gas (Xe; teal spheres). Phenylalanine 229 (F229; cyan sticks) forms a wall of the hydrophobic CO alcove (green sticks) and is proposed to facilitate the Ni_p_-CO bond. PDB 2Z8Y.

The EPR and Electron Nuclear Double Resonance (ENDOR) spectroscopic studies described here reveal that the A-cluster binds not one but two CO. It also shows that substituting the highly conserved residue Phe229, which forms one of the walls of the alcove, with the smaller Ala significantly destabilizes CO binding to the A-cluster and labilizes the conformation of the CO-bound cluster, whereas the rather conservative substitution F229W causes much smaller perturbations. This paper thus demonstrates the key role of an intact alcove in forming and stabilizing the native conformation of the Ni(I)-CO intermediate in the Wood-Ljungdahl pathway of anaerobic CO and CO_2_ fixation.

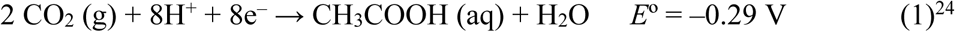

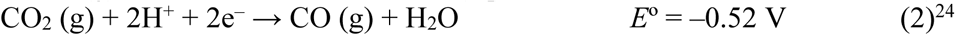

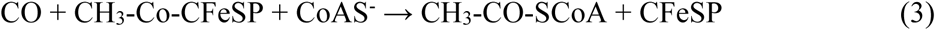

## Materials and Methods

### Generation, Growth, Lysis, and Purification of ACS Alcove Variants

ACS F229 variants were prepared from the *acsB* gene, representing the Acetyl-CoA Synthase subunit of CODH/ACS and was placed into a *lac*-inducible, kanamycin-resistant, and C-terminal His6-tagged pET29a vector and known as pET29ACSMT_HT_. ACS variants were generated via site-directed mutagenesis using a QuikChange II kit from Agilent Technologies. While the wildtype was left unadulterated, primers (Integrated DNA Technologies) used in generating the F229A and F229W variants contained the following sequences: F229A – *sense* 5’-G CGG GCT GGT ATG ATG <u>GCC</u> GGT GGC -3’ *antisense* 5’-GGT AAC GCC ACC <u>GGC</u> CAT CAT ACC A -3’; F229W – *sense* 5’-C CTG CGG GCT GGT ATG ATG <u>TGG</u> GGT -3’ *antisense* 5’-GT AAC GCC ACC <u>CCA</u> CAT CAT ACC AGC -3’;. The plasmids were co-transformed into competent BL21(DE3) *E. coli* with the genes *iscS-iscU-iscA-hscB-hscA-fdx* containing [4Fe-4S] construction machinery from *A. vinlandii* within the ampicillin-resistant, *ara*-inducible pBD1282 plasmid for proper [4Fe-4S] construction.^25^ BL21(DE3) cells harboring the ACS variant plasmids were grown as described previously.^25^ Cells harboring the ACS variant plasmids were lysed and purified as described previously, with concentrations measured by Rose Bengal protein concentration assay.^26^ Buffers and reagents used in purification were anaerobically prepared (≤ 2.5 ppm O_2_), and all glassware was acid washed for at least one hour prior to usage in order to remove contaminating metal.

### ACS Metal Reconstitution

ACS A-cluster reconstitution was performed as previously described.^27^ Metal content was assessed through ICP-OES at the University of Georgia’s (Athens, GA) Center for Applied Isotope Studies (CAIS) and compared to total ACS concentration verified by Rose Bengal assay.^26^

### Titanium (III) Citrate Reductant Preparation

Anaerobic solutions of 83 mM titanium (III) citrate reductant were prepared as described previously.^28^ To prevent excess photodegradation, Ti(III) citrate solutions were stored in brown glass away from ambient light, and were stored at room temperature under anaerobic conditions.

### Preparation of ENDOR samples

ENDOR samples of ACS variants were anaerobically prepared to 300 µM in the presence of 2 mM sodium dithionite (Sigma) in a sealed and crimped 2 mL vial, where the vial headspace was purged with either 100% ^12^CO gas (Cryogenic Gases; Detroit, MI) for 20 minutes or 100% ^13^CO gas (Cambridge Isotope Laboratories; Tewksbury, MA) for 10 minutes. Once the sample was purged with CO, it was transferred via syringe to a quartz Q-band EPR tube (2 mm diameter) placed inside of a rubber septum-sealed X-band EPR tube and a headspace either 100% ^12^CO gas or 100% ^13^CO gas for the respective isotope. Once the ACS variant sample was properly placed within the inner Q-band tube by syringe, the CO-containing rubber septum-sealed X-band tube was removed from the anaerobic chamber and immediately frozen in liquid nitrogen. Once frozen, the Q-band tube was removed from the X-band tube and cryogenically stored for subsequent spectral acquisition.

#### EPR and ENDOR Spectroscopy

35GHz continuous wave (CW) EPR and ENDOR were collected on a previously described spectrometer that is equipped with a Janus liquid helium immersion dewar for measurements at 2K.^29-31^ These CW measurements employed 100 kHz field modulation, with experiments performed in dispersion mode under rapid passage conditions, which yields an absorption-display spectrum. Derivative-display spectra were generated numerically by applying a Savistky-Golay filter to the 35 GHz field-swept absorption experimental spectra in LabCalc.

For a single molecular orientation of a nucleus with spin *I* = 1/2 (^13^C, ^1^H), the ENDOR transitions are described in first-order described by the equation:

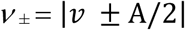

where *v*_n_ is the nuclear Larmor frequency and A is the orientation-dependent hyperfine coupling When *v*_n_ > A/2, as in all spectra reported here, the ENDOR pattern is a doublet split by A, and centered at *v*_n_.

All EPR, and ENDOR simulations were obtained using the EasySpin simulation package. The fits of the EPR spectra were obtained using the *esfit* function in EasySpin.^32^

## Results

### EPR Spectra of A_red_-^13^CO in Wild Type (WT) and in F229W and F229A Variants

**Figure 2** shows the absorption-display (black) and numerical derivative (red) Q-band EPR spectra of the WT A_red_-^13^CO and corresponding spectra of this intermediate for the F229A and F229W mutants, collected at 2K under dispersion rapid-passage conditions. The Q-band spectrum of WT A_red_-^13^CO exhibits a signal describable by an axial g-tensor, g_⊥_ = 2.085 and g_‖_ =2.036.^33^ The F229W A_red_-^13^CO mutant shows a similar Q-band spectrum, but the center is somewhat perturbed, with g_⊥_ = 2.081 and g_‖_ =2.037, **Table 1**. The g-tensors of these signals are long known to be characteristic of Ni(I) with a d_z_^2^-based odd-electron orbital.^34^ The similarity of the spectra for WT and F229W enzymes indicates that the A_red_-^13^CO state of the variant retains the WT conformation, largely unperturbed by the relatively conservative replacement of F229 by W. However, the enhanced resolution at Q-band shows slight mutation-induced shifts in g-values, **Table 1**, indicative that replacement of the F residue with the larger (and potentially H-bonding) W residue causes some distortion to the metal center. More intriguingly, we also note the extremely weak intensity in the F229W absorption-display spectrum (highlighted by a box; suppressed in the derivative display) at fields above the high-field edge of the WT spectrum (above ∼12,300 G). We return to this feature, below.

**Table 1.** g-values measured at Q-band for *A*_*red*_*-CO* as formed in ACS variants F229 (WT), F229W, and individual conformers of F229A. The g-values and percentage populations for the F229A conformers were obtained by a decomposition of the EPR spectrum as described in text and **SI**.

| | $g_{\perp}$ | $g_{\parallel}$ | |
| --- | --- | --- | --- |
| F229 (WT) | 2.085 | 2.036 |  |
| F229W | 2.081 | 2.037 |  |
| F229A |  |  | Population (%) |
| WT-Conformer | 2.078 | 2.031 | 20 |
| A-Conformer | 2.017 | 2.065 | 60 |
| O-Conformer | 2.055, 2.048 | 2.021 | 20 |

**Figure 2.**
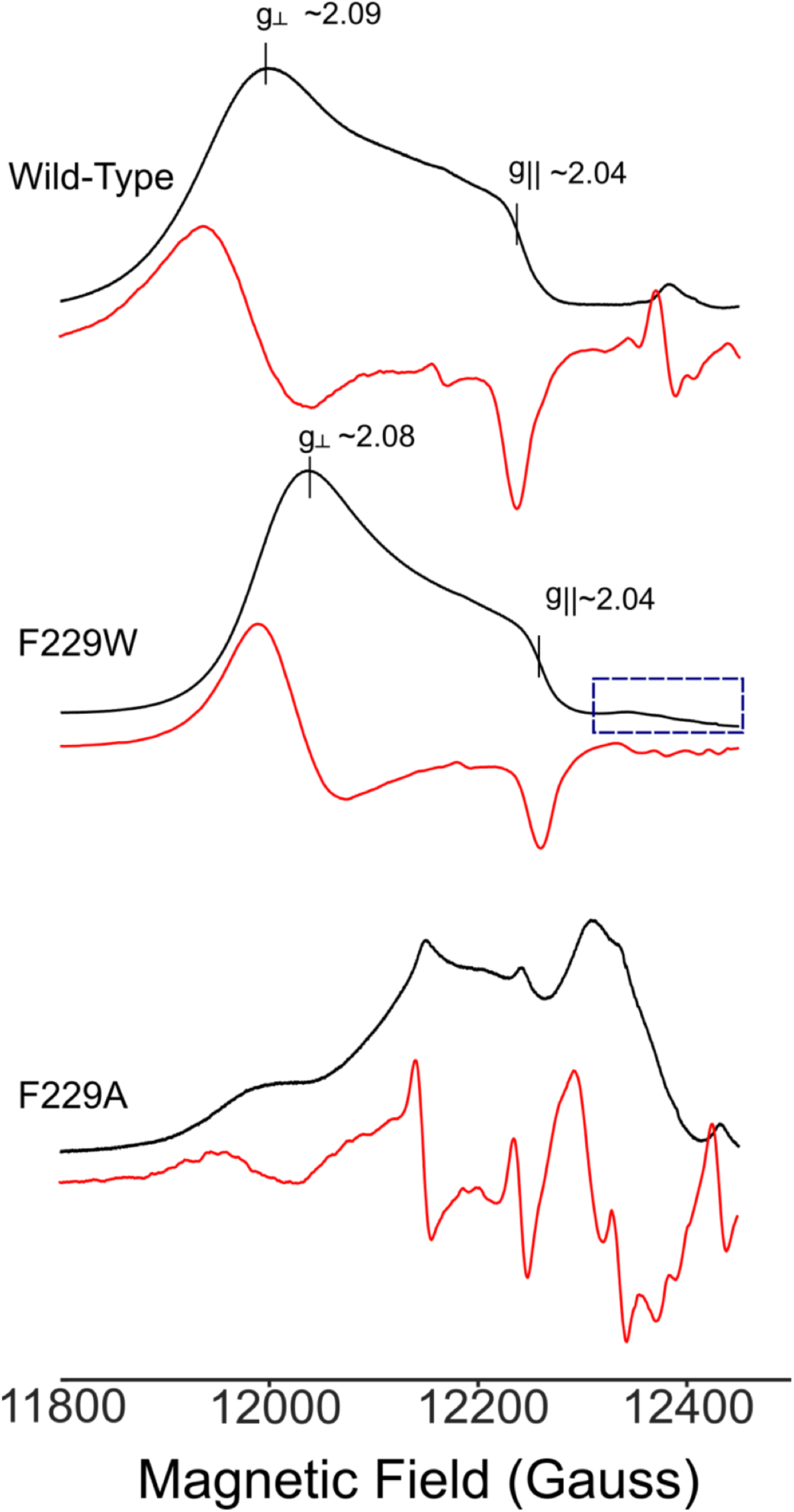
35 GHz CW rapid-passage absorption-display EPR spectra (Black), numerical derivatives (red) of *A*_red_-^13^CO. (*Top*) wild-type; (*Middle*) F229W mutant (dashed box is a weak response from 2^nd^ conformer, see text); (*Bottom*) F229A; notice how weak but sharp features are enhanced in the derivatives. *Experimental conditions*: Microwave frequency ∼34.9GHz, microwave power, 100mW; modulation amplitude 1G, time constant = 32ms; temperature 2K.

In contrast, the A_red_-CO center is strongly perturbed by the replacement of F229 with the much smaller alanine residue. This is seen firstly in the EPR signal intensity of the CO-treated F229A variant, which is about 10-fold lower than that of WT or the F229W variant, but more dramatically by the sharply different EPR spectrum of F229A-A_red_-^13^CO, which has additional features not observed in the WT NiFeC signal (**Figure 2**). There is no direct way to decompose this spectrum into its contributing components, but as will be shown below, it can be decomposed into three contributions by reference to ^13^C ENDOR measurements of this species. However, this variant binds CO poorly: the combined intensities of the different species amount to only about 10% of those observed for WT and F229W.

### ^13^C ENDOR of the A_red_-^13^CO states of WT enzyme and F229 Variants

***A***_***red***_***-***^***13***^***CO(WT):* Figure 3** shows a 2D field-frequency pattern of ^13^C ENDOR spectra collected across the EPR envelope of WT A_red_-^13^CO. The correspondence of ENDOR spectra to the fields of observation within the envelope is shown through inclusion of the 35 GHz CW rapid-passage EPR spectrum on the left of the figure. The most pronounced feature in the ^13^C ENDOR spectra is the peak whose offset from the ^13^C Larmor frequency (*v*_C_) by (*v*-*v*_C_) ∼ 15-16 MHz, and whose intensity roughly tracks the intensity of the A_red_-^13^CO(WT) EPR spectrum. This peak, denoted C1, is assigned as the *v*_+_ signal of ^13^CO axially bound to the d_z_^2^ SOMO of Ni_p_. Its frequency is essentially field independent and corresponds to an isotropic ^13^C hyperfine coupling, a_*iso*_(C1) *∼* 31 MHz. However, the C1 ENDOR peaks are relatively broad and show variations in linewidth across the EPR envelope, indicative of unresolved dipolar hyperfine interactions with maximum value of ∼ 5-6 MHz. It is well understood why a metal-bound ^13^CO should have such a disparity in the magnitudes of the isotropic and anisotropic hyperfine interactions. For spin density in an *sp* orbital on the ^13^C of CO sigma-bonded to a metal ion, one expects as the ratio of the maximum ^13^C dipolar coupling to the isotropic coupling, a value, a_*iso*_ /2T*∼* 20/1,^35^ and the total breadth is roughly as anticipated for the sum of the dipolar couplings from spin on ^13^C and on the Ni.

**Figure 3.**
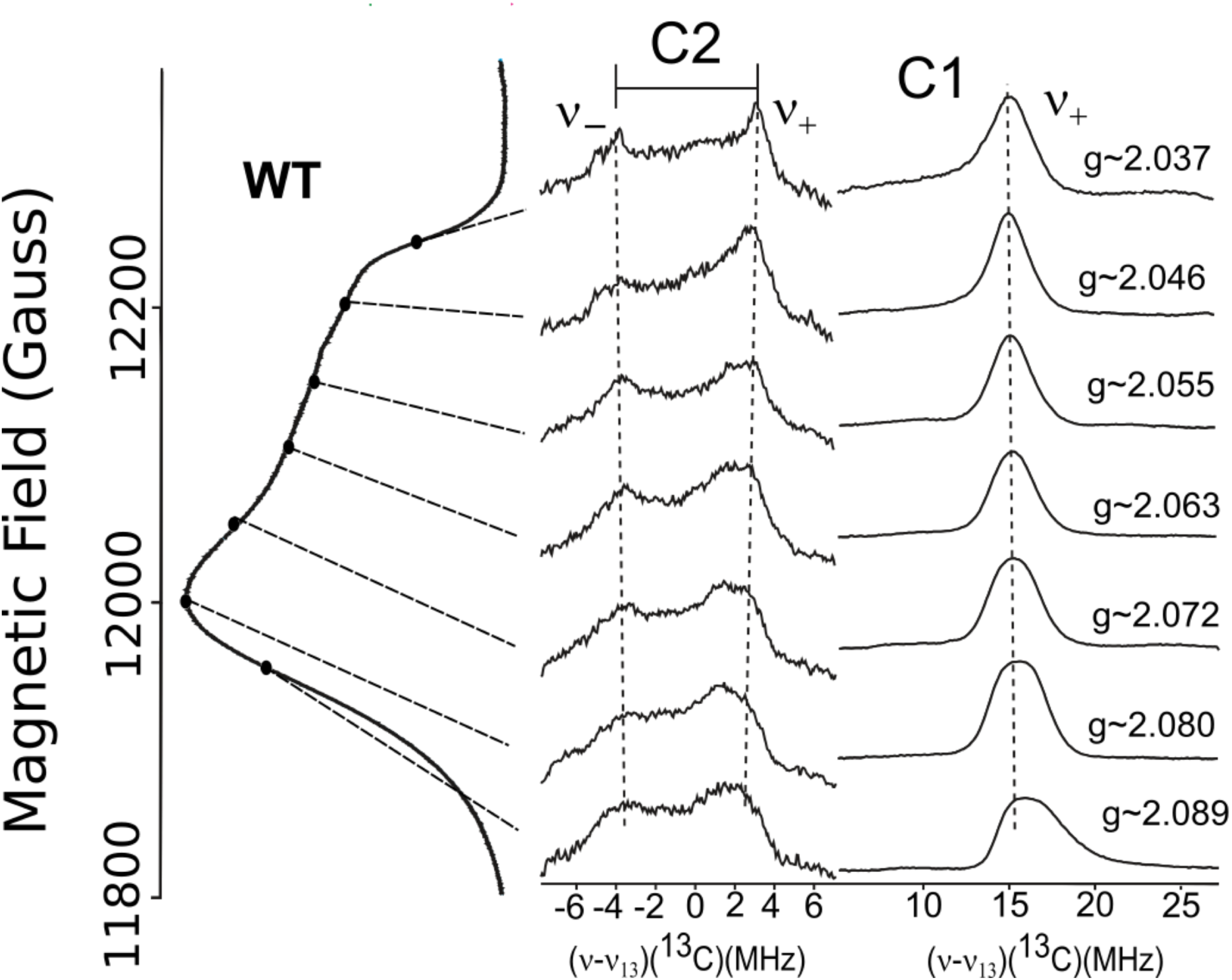
35GHz 2D field-frequency pattern of ^13^C CW ENDOR spectra (center, right), keyed to rapid-passage EPR spectrum, of A_red_-^13^CO wild-type (WT) (left). *ENDOR experimental conditions:* Microwave frequency ∼34.9GHz, modulation amplitude 1G, time constant = 64ms, RF sweep rate 0.5MHz/s; number of scans = 20; temperature 2K. Both center, right panels show frequency relative to the ^13^C Larmor frequency, 13 MHz; center panel amplitude x2.

Surprisingly, there is a signal from a second ^13^C (labeled, C2) centered at the ^13^C Larmor frequency, *v*_C_, that also tracks the A_red_-^13^CO(WT) EPR spectrum. As the field is varied, this signal changes in shape and in total breadth. The maximum breadth, which corresponds to a maximum hyperfine coupling, occurs near g_‖_, where the spectrum shows a resolved doublet corresponding to a *v*_+_/*v*_-_ pair split by A *∼* 6-7 MHz. Although the poor resolution of this signal implies a site with a considerable distribution in binding geometries, simulation of the 2D field-frequency pattern is instructive. In the best simulations (**See Supplementary Information**) the pattern is satisfactorily described by a largely isotropic hyperfine interaction: *a*_*iso*_ ∼ 5.2 MHz and maximum dipolar coupling of ∼ 1.5 MHz, but with a spread in the couplings subsumed in the imposition of a large ENDOR linewidth.

How is this second ^13^CO bound? Its properties are most plausibly explained if it is bound in the Ni_p_ ‘equatorial plane’, namely roughly normal to the axis of the d_z_^2^ odd-electron orbital, where its isotropic coupling is small because of greatly diminished overlap with the d_z_^2^ SOMO. **Figure 4** gives a plausible illustration of such a structure. One might instead suggest that the second CO binds, again ‘on axis’, to Ni_d_, with ∼1/5^th^ of the spin density on Ni delocalized to its d_z_^2^ orbital, which would introduce an isotropic hyperfine coupling of the observed magnitude this axial ^13^CO. However, given the large distance between the two Ni, there can be no direct overlap between the two d_z_^2^ orbitals, and any such spin on Ni_d_ would necessarily arise via polarization of the Ni-S bonds to the bridging sulfurs, and it is implausible that such a process could generate that high a spin density in the d_z_^2^ orbital of Ni_d_. Thus, we propose that the structure of A_red_-^13^CO(WT) is as visualized in **Fig 4**.

**Figure 4.**
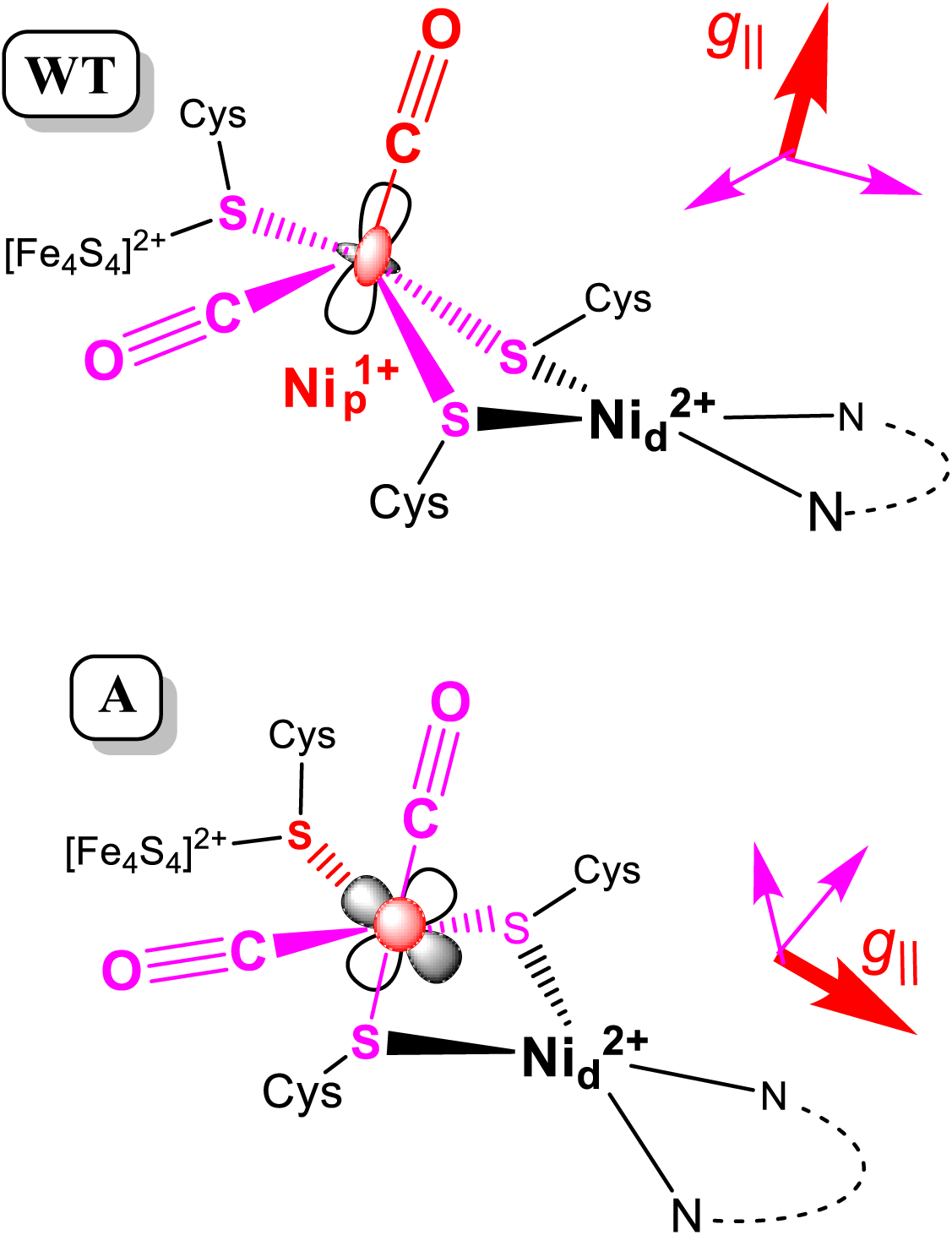
Sketch of proposed structures for WT and A conformers of A_red_-CO. Axes and bonds/atoms: *Red*, along g_‖_ ; *Purple*, associated with the g_⊥_ plane.

### A_red_-^13^CO(F229A)

**Figure 5** shows the ^13^C CW 2D field-frequency ENDOR pattern of F229A A_red_-^13^CO accompanied by the 35 GHz CW absorption-display EPR spectrum. The intensity of the NiFeC signal from this variant is approximately ten-fold lower than from WT or the F229W variant, demonstrating that disruption of the alcove through introduction of the smaller alanine residue destabilizes reduction and CO binding to the A-cluster. In addition, the F229A substitution introduces extensive cluster flexibility. Thus, the experimental EPR spectrum is overlaid by the spectrum from a fit that decomposes it into three distinct contributing components: i) a conformer denoted, A, with g_‖_ > g_⊥_ ∼ 2; ii) a conformer equivalent to that of the wild-type with g_⊥_ > g_‖_ ∼2, and thus denoted WT; iii) a component termed O, also with g_⊥_ > g_‖_ ∼2, but with g_⊥_(O) < g_⊥_ (WT). The contributing spectra of these components also are shown; the g-tensors, linewidths, and fractional contributions of the three components are listed in **Table 1**. As now described, this decomposition was guided by analysis of the 2D pattern of ^13^C ENDOR measurements displayed in the figure.

**Figure 5.**
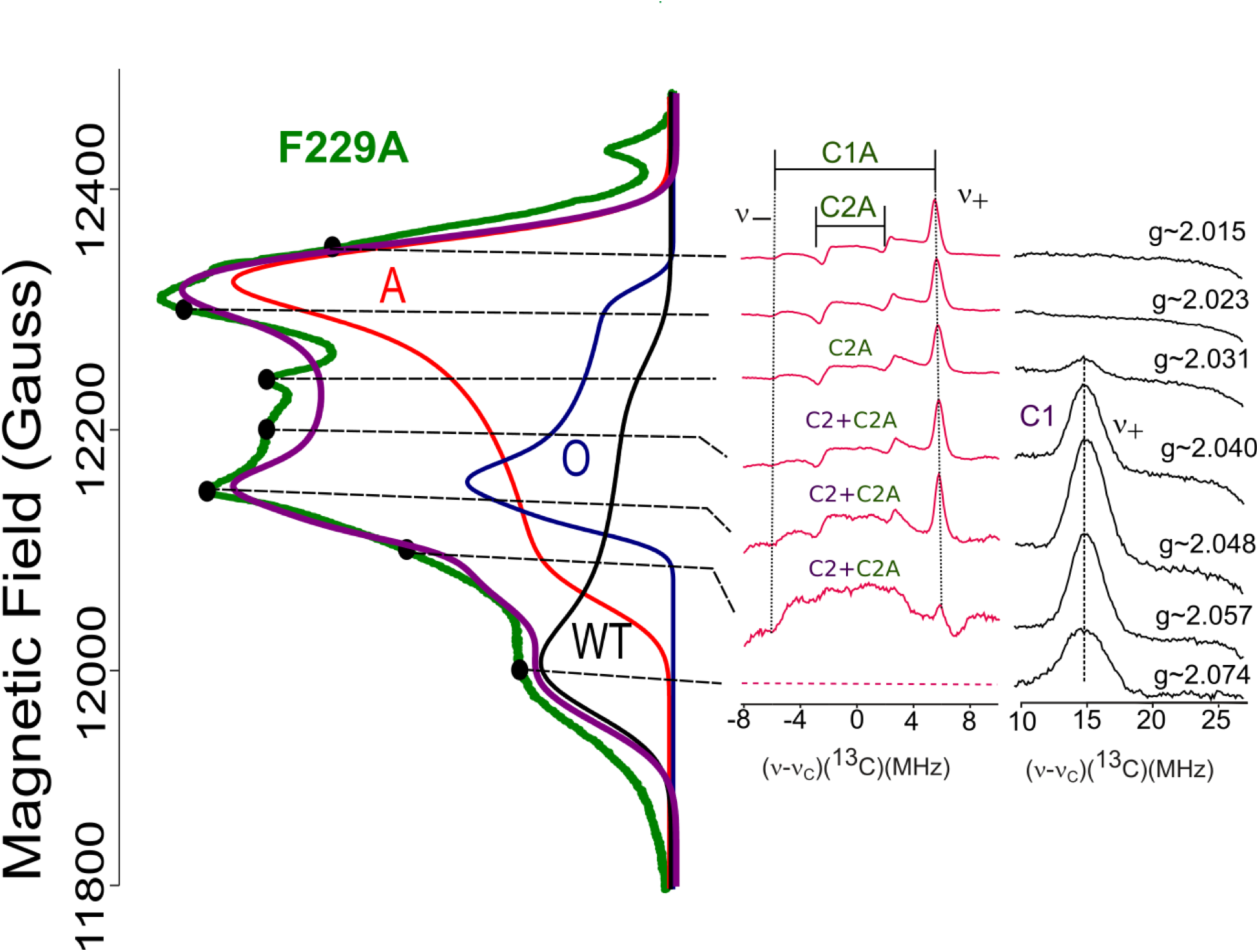
35 GHz 2D field-frequency pattern of ^13^C ENDOR spectra (center, right) keyed to CW rapid-passage EPR spectrum (left) of A_red_-^13^CO-F229W (left). EPR spectrum is overlaid with a sum simulation with contributions from three species: 1) **W** - wild-type conformer,. **g** = [2.07825 2.031]; 2) **O** – previously reported; **g**= [2.0555 2.048 2.021]; 3) **A** –component; g = [2.0169 2.065]. As described in Materials & Methods and SI, simulation and decompositions obtained using the *esfit* function in EasySpin, by varying the linewidths, g-values, and relative weights of each component. The fit was achieved using a generic algorithm (RMSD = 0.034). Both center, right panels show frequency relative to the ^13^C Larmor frequency, 13 MHz. *ENDOR experimental conditions* as in **Fig 2**.

Firstly, the low-field shoulder of the EPR spectrum is at ∼ 12 kG, corresponding to g_⊥_ (WT), and is identified with the WT conformer by the ENDOR observation of the *v*_+_ feature from ^13^C1 of the WT enzyme (A_*iso*_ *∼* 31 MHz), which has a field-dependent intensity that roughly parallels the intensity of the EPR signal from the WT conformer. A second, sharper signal at (*v*-*v*_C_) ∼ 5-6 MHz, denoted C1A, again shows an isotropic coupling (a_*iso*_(C1A) ∼ 11 MHz) but it is most intense at the high-field edge of the EPR spectrum (∼12,350 G), where there is no WT EPR response, and it decreases in intensity as the field decreases and vanishing below ∼ 12,000 G. The high-field, high-intensity edge of this ENDOR pattern is associated with the EPR intensity maximum at that field, which we ascribe to g_⊥_ of a ^13^CO-bound conformer, denoted A, having g_⊥_ ∼ 2.02 < g_‖_ ∼ 2.07, with the value for g_‖_ suggested by the vanishing of the C1A ENDOR signal at low field. The C1A signal is therefore assigned as the *v*_+_ feature of ^13^CO bound in the A conformer.

In addition, a doublet of ENDOR peaks centered at the *v*_13C_, denoted C2A, also tracks the A-conformer EPR signal. This doublet corresponds to the *v*_+_/*v*_-_ partners of a second bound ^13^CO, which has a smaller and clearly isotropic coupling, a_*iso*_(C2A) ∼ 4.8 MHz. Where the A and WT conformer EPR signals overlap, the C2A doublet from the A conformer overlaps with the C2 doublet associated with the WT conformer, and resolution is lost. However, the sharp lines of the isolated C2A signal imply a well-defined binding geometry. Parallel considerations to those raised in discussion of the origin of the C2 signal, when applied to the sharp C2A signal, clearly indicate a covalently-bound Ni ligand, and not a non-covalently bound CO trapped in the collapsed alcove. This in turn supports our preference for assigning the WT C2 signal to a covalently bound ^13^CO.

The other prominent maximum in the EPR spectrum, at ∼ 12,150 G, is not accounted for by contributions from either the WT or A conformers, and is assigned as the g_⊥_ feature of yet another conformer, denoted O. The O conformer again has (g_⊥_ > g_‖_ ∼ 2) and thus a d_z_^2^ SOMO, as in the WT. However, the absence of any corresponding ^13^C signal means the O conformer does not have any bound ^13^CO.

With this guidance, the observed EPR spectrum for this state was fit to the three component spectra presented in **Fig 5**, as described in M&M, with output g-value and fractions listed in **Table 1**. As can be seen, the fit accounts for all significant features in the EPR spectrum. The g tensor and the two ^13^C hyperfine couplings of the WT component of the F229A intermediate match those of WT enzyme itself, indicating that this conformer indeed has the structure seen in WT enzyme, basically unperturbed, with Ni_p_ having a d_z_2 SOMO and binding both axial and equatorial ^13^CO. The g-tensor of the A conformer is like that of Cu(II), and instead indicates the Ni_p_ SOMO is a ‘cloverleaf’ orbital (d_x_^2^_-y_^2^, d_xz_, etc). Interpretation of the ENDOR measurements is a challenge. Were these two ^13^CO bound in the plane of and along the lobes of a cloverleaf orbital, the strong overlap would yield large isotropic couplings, of the same order as those of C1 of WT. The small couplings require that an in-plane ^13^CO would need to bind roughly along the cloverleaf nodes.

### A_red_-^13^CO(F229W)

**Figure 6** shows the 35 GHz ^13^C CW 2D field-frequency ENDOR pattern for F229W A_red_-^13^CO, accompanied by its CW 2K rapid-passage EPR spectrum. This state also shows a strong *v*_+_ ^13^C1 ENDOR signal that tracks the dominant WT component of its EPR signal, disappearing at higher and lower field than this signal. This ENDOR response again shows the isotropic coupling (a_*iso*_(C1) *∼*31 MHz) of a ^13^CO bound on the z-axis of the d_z_^2^ orbital of Ni_p_, as in the WT intermediate. Likewise, there is also the poorly-resolved C2 intensity that indicates the presence of the second ^13^CO seen with the WT intermediate; it is referred to again in the next paragraph. The EPR and ENDOR thus show that the metal center of A_red_-^13^CO(F229W) is predominantly in the WT conformation, largely unperturbed from that of the WT intermediate, with a d_z_^2^ odd-electron orbital and two bound ^13^CO, one ‘out-of-plane’ (C1), the other likely bound ‘in-plane’ (C2).

However, there is an additional ^13^C ENDOR signal in the 2D pattern of A_red_-^13^CO(F229W) at (*v*-*v*_C_) ∼ 6 MHz (denoted C1A), along with a doublet centered at *v*_C_, denoted C2A, both of whose intensities maximize at fields above the high-field edge of the WT EPR spectrum and track the EPR spectrum derived above for the A-conformer seen in the F229A intermediate (**Fig 7)**. The C2A signals overlap with the C2 signal from the WT conformer at fields within its EPR spectrum, which causes the poor resolution of the ENDOR signals within ∼ ± 4 MHz of *v*_C_. Thus, it is firstly clear that the F → W conversion leaves a dominant, unperturbed WT conformer with a d_z_^2^ odd-electron orbital on Ni_p_ and with both an axial (C1) and an equatorial (C2) bound ^13^CO. Secondly, however, the change in residue also induces the formation of a small percentage of the A conformer found in the F229A variant, in which Ni_p_ exhibits a cloverleaf (eg. d_x_^2^_-y_^2^) odd electron orbital and with two ^13^CO likely bound in-plane.

**Figure 6.**
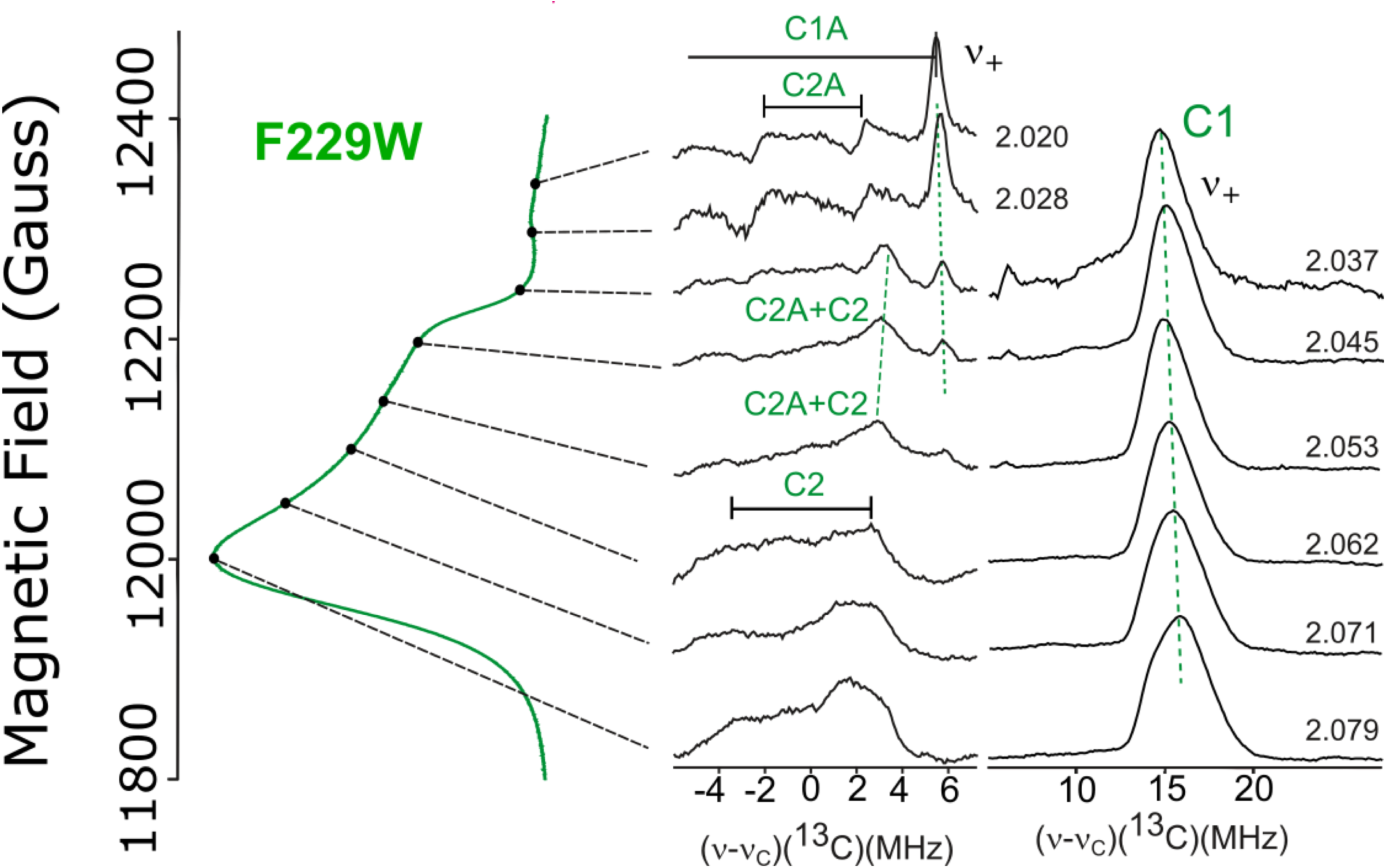
35 GHz 2D field-frequency pattern of ^13^C ENDOR spectra (center, right) keyed to CW rapid-passage EPR spectrum (left) of A_red_-^13^CO F229W. Both center, right panels show frequency relative to the ^13^C Larmor frequency, 13 MHz *ENDOR experimental conditions:* As in **Fig 2**.

**Fig 7.**
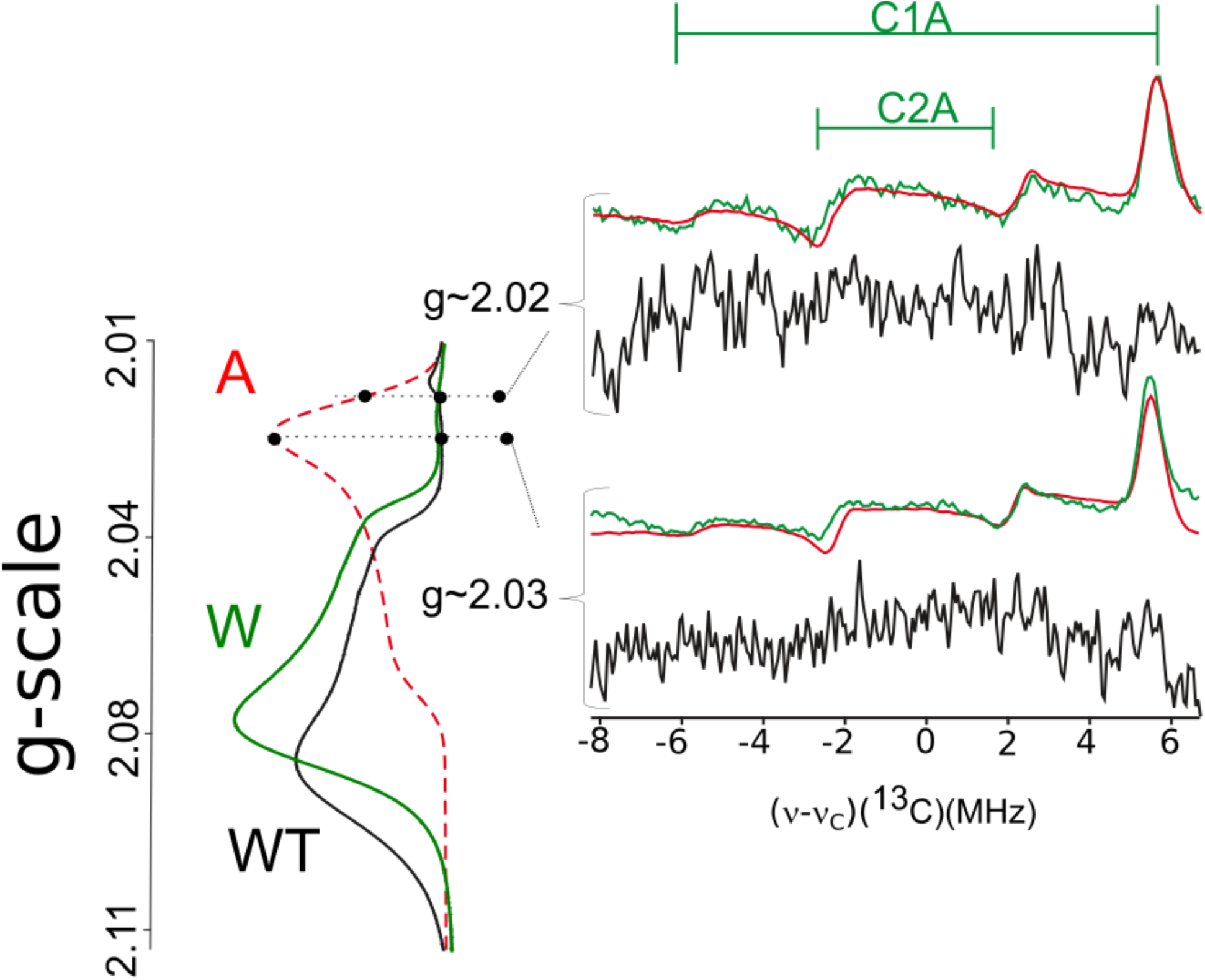
35 GHz ^13^C CW 2D field-frequency ENDOR spectra keyed to 35GHz CW rapid-passage EPR spectra of WT (black) and F229W (green) *A*_red_-^13^CO. The experimental EPR spectra are overlaid with a simulation of the **A** component of the decomposition of the F229A mutant spectrum (red dashed). All ENDOR data shows frequency relative to the ^13^C Larmor frequency, 13 MHz *ENDOR* e*xperimental conditions*: as in **Fig 2**

## Discussion

Previously, we were only aware of a single CO binding in the WT A_red_-CO state of ACS, as definitively shown by X-ray absorption spectroscopy.^21^ Here 35 GHz EPR and ENDOR studies showed that the WT Ni_p_(I)-CO conformation exhibits a d_z_^2^ odd-electron orbital with CO bound on the orbital axis. X-ray crystallography revealed a gas-binding ‘alcove’ at the end of the hydrophobic tunnel from CODH to the active site of ACS, containing Xe as a CO surrogate, 3.9 Å above the A-cluster proximal Ni_p_, (See **Fig 1**).^5^ The highly conserved residue Phe229 forms one of the walls of the alcove, and thus was implicated in delivering CO to the ACS active site. We further suggest that this axial Ni_p_(I)-bound CO occupies and is ‘cradled’ by the alcove when bound in the A_red_-^13^CO(WT) intermediate.

The present EPR and ENDOR studies also revealed a second bound CO, likely coordinated in the equatorial plane of the Ni_p_ d_z_^2^ orbital. The binding of a second CO to the A-cluster during sample preparation under a high partial pressure of CO could explain the inhibition of the CoA/acetyl-CoA exchange (K_i_ = K_d_ = 0.4 mM)^36^ and acetyl-CoA synthesis (50% inhibition at ∼ 0.45 mM)^37^ activities of ACS at elevated CO concentrations. In stopped-flow FTIR experiments, a dominant peak (1997 cm^-1^) appeared at about the same rate as formation of the NiFeC species; at a slower rate and much lower intensity, a second peak appeared at 2044 cm^-1^, which probably corresponds to the more weakly coupled ‘in-plane’ CO.^12^

The finding of this second bound CO speaks to the conformational preferences of ACS, which has been viewed through the lens of a two-state open/closed model. A recent CODH-ACS crystal structure reveals a snapshot of the CO-bound form, which adopts the closed conformation. It contains a single axial CO bound to Ni_p_ (Cohen et al, 2020, submitted), with Phe229 and Ile146 positioned near the A-cluster to promote formation of the CO-bound state. In this closed state, Ile146 appears to block the site where we suggest the second CO might bind. However, in the “open” ACS conformation, these residues undergo a major reorientation, which favors methylation. There is strong evidence for a “two-substrate-bound” state in which both methyl and CO are independently coordinated to Ni_p_ but have not yet undergone C-C bond formation.^38^ There are no structures of this state, which we predict to be transient because it would quickly form acetyl-Ni_p_. However, in support of this idea, the recent CO-bound crystal structure and a cryo-EM analysis (personal communication, Cohen and Drennan) reveal ACS possesses much more flexibility than formerly appreciated. This plasticity is the means to accommodate the two-substrate-bound state. We propose that the state revealed by the present work to contain two CO bound to Ni_p_ is a surrogate for and provides a first glimpse of this important intermediate in the catalytic mechanism.

The present studies further show that substituting Phe229 with the smaller Ala significantly destabilizes CO binding to the A-cluster and labilizes the conformation of the CO-bound cluster, while the rather conservative substitution F229W causes much smaller perturbations. Thus, by constraining the location of the Ni_p_-bound CO, an intact alcove helps stabilize the WT conformation of the ACS metallo-center, a role that evokes the idea the native conformation of an enzyme can generate an ‘entatic state’ in which the metal center is activated for catalysis.^39-40^

The dominant form of the A_red_-CO state of the F229W variant shares its conformation and electronic structure (g_⊥_ > g_‖_ ∼ 2) with WT, but a small minority also adopts an “A” conformation in which Ni_p_ exhibits the reversed g-tensor (g_‖_ > g_⊥_ ∼ 2) characteristic of a cloverleaf (d_x_^2^_-y_^2^-like) odd-electron orbital. This conformer also binds two CO, but with distinctly smaller hyperfine interactions. In contrast to these two enzyme variants, substitution of the phenyl sidechain of F229 with the three-fold smaller residue of Ala, disrupts the alcove and in doing so introduces a conformational flexibility to the A_red_-CO state, allowing rearrangements that highly perturb the NiFeC center. The F229A variant not only exhibits significantly diminished formation of the Ni(I)-containing A_red_-CO than WT or the F229W variant, but the substitution introduces a high degree of cofactor flexibility. The Ni(I) F229A enzyme is mostly (60%) in the “A” conformation with Ni_p_(I) binding two in-plane CO. In addition, this variant exhibits equally small amounts (20% each) of residual WT conformer in which Ni_p_(I) binds one axial CO and another in plane, and of a novel “O” state that does not bind CO at all. Considering the latter, previous attempts to reduce the Ni_p_(II) A-cluster to the A_red_ state in the absence of CO have been fruitless, even using strong chemical reductants or highly negative electrochemical potentials. To explain the formation of the Ni(I) “O” state, we speculate that disruption of the alcove by the F229A mutation allows formation of the “closed channel-open” conformation of ACS, primed for reductive carbonylation,^5^ while removal of an alcove wall also permits CO escape, resulting in formation of only a small proportion of the CO-free A_red_ O state. The present results thus both reveal the binding of two CO, not just one, to the A center of ACS and highlight the key role of an intact alcove in formation and stabilization of the Ni(I)-CO intermediate in the Wood-Ljungdahl pathway of anaerobic CO and CO_2_ fixation.

## Acknowledgements

This work was supported by the NSF (MCB-1908587, BMH) and the NIH (R37 GM039451, SWR). We are grateful to Drs. Catherine Drennan, Seth Cohen and Ritimukta Sarangi for discussions of the structural implications of these ^13^C-ENDOR results.

---

TOC Figure

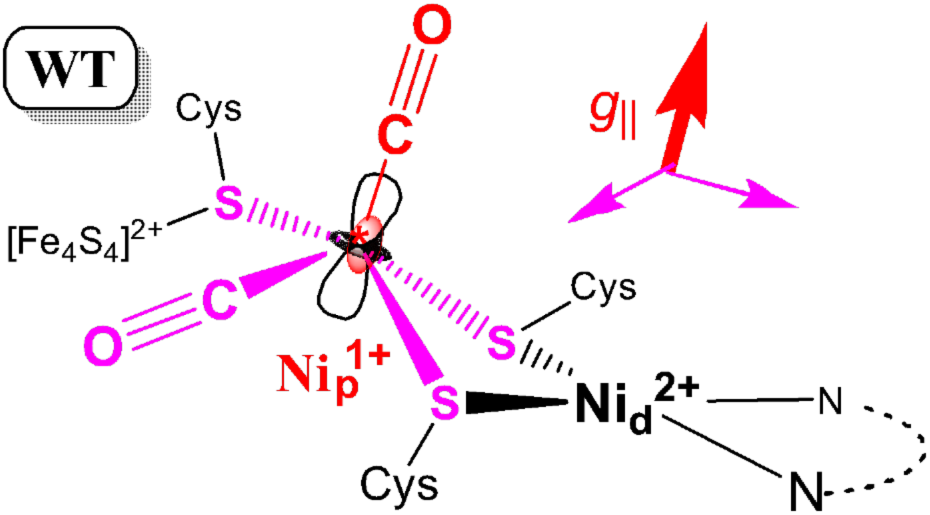

**Scheme 1.**
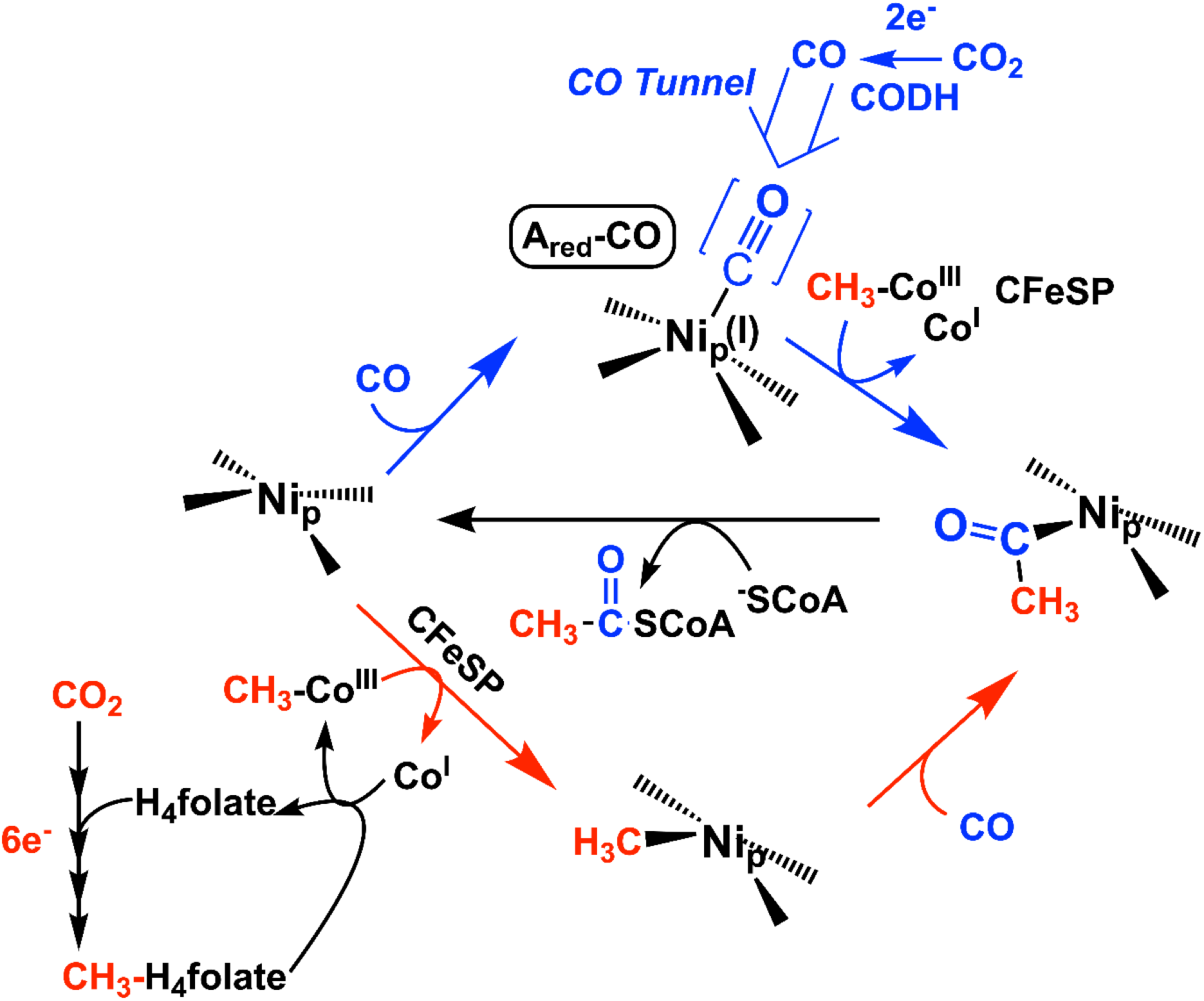
Paramagnetic mechanism for ACS catalysis. See text for details.

## Notes

### Competing Interest Statement

The authors have declared no competing interest.

