## Supplementary Information for "^13^C Electron Nuclear Double Resonance Spectroscopy Shows Acetyl-CoA Synthase Binds Two Substrate CO in Multiple Binding Modes and Reveals the Importance of a CO-Binding ‘Alcove’"

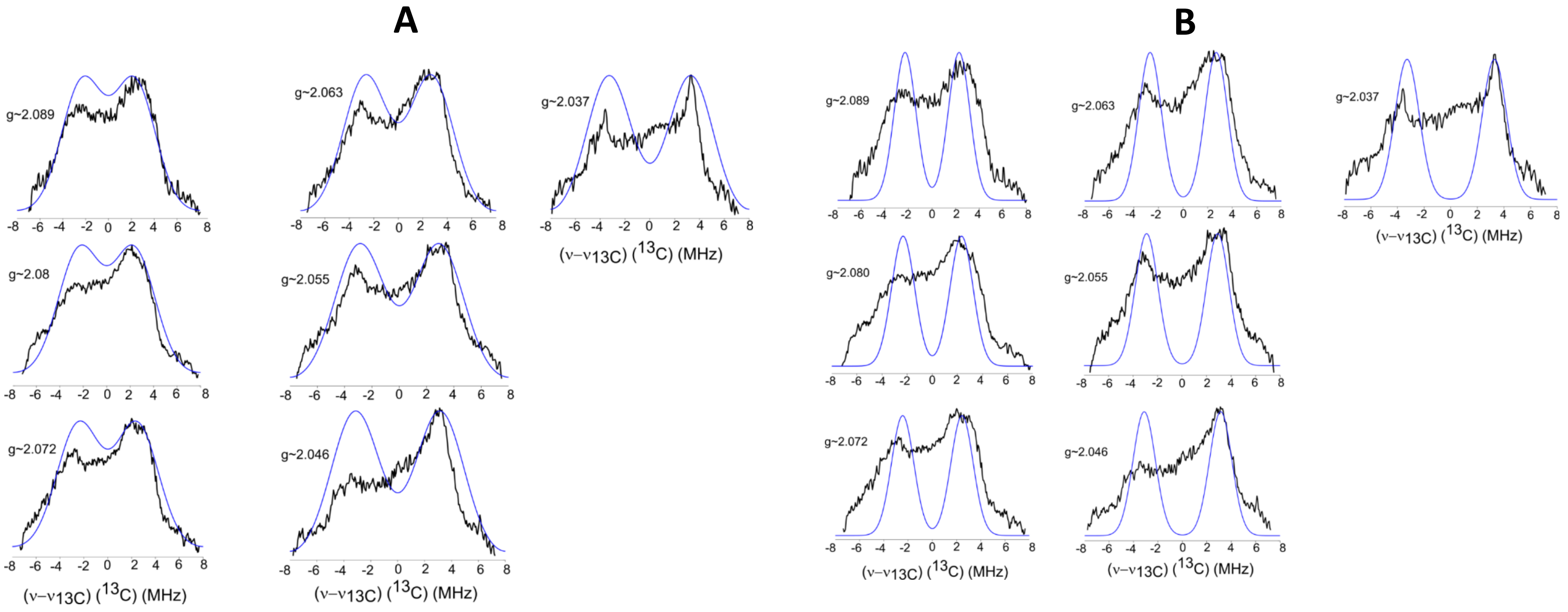


**Figure S1.** ENDOR spectra (black) overlaid with simulations (blue) of the $^{13}$C2 signal from $A_{red}$-$^{13}$CO (WT); frequencies relative to $^{13}$C Larmor frequency, ν($^{13}$C). Simulations were performed using matrix-diagonalization algorithm in the *salt* function of the EasySpin simulation package, yielding an axial A-tensor, coaxial with **g**: $A_{\perp}$ = 4.53, $A_{\|}$ = 6.71 MHz. ***Panel A***: ENDOR linewidth ENDOR linewidth, 4.25 MHz. ***Panel B***: linewidth decreased to 2.15 MHz to better visualize the field dependence of peak frequencies. *ENDOR experimental conditions:* Microwave frequency ~34.9GHz, modulation amplitude 1G, time constant = 64ms, RF sweep rate 0.5MHz/s; number of scans = 20; temperature 2K

**Fig S2**: EasySpin script for fitting (**A**) and simulating (**B**) the F229A $A_{red}$-$^{13}CO$ EPR spectrum.

**S2A.** *esfit* EasySpin script used to **fit** the components of the F229A $A_{red}$-$^{13}CO$ EPR spectrum. The fitting protocol used the *pepper* function in Easyspin, and varied the g-values, linewidths, and relative weights (total weight = 100%) for the individual components of the F229A $A_{red}$-$^{13}CO$. Output of fits listed in **Table 1** of main text

```
clc; clear all;
%calls data
[B,spc] = textread('1907152.txt','%f %f'); %reads in experimental data for F229A

%Component 1
Sys1.g = [2.0555 2.048 2.021]; %g-values for 1st component
Sys1.lw = 2.85024;
Sys1.weight = 0.203;

%Component 2
Sys2.g = [2.0169 2.065];
Sys2.lw = 5.75413;
Sys2.weight = 0.597;

%Component 3
Sys3.g = [2.07825 2.031];
Sys3.lw = 7.775;
Sys3.weight = 0.2;

Exp.mwFreq = 34.85;     % GHz
Exp.Range = [1150 1248];% mT
Exp.Harmonic = 0; %absorption spectrum

Exp.Temperature = 2; % 2K CW


Opt.Method = 'perturb';
Opt.nKnots = [61 0];
Exp.Harmonic = 0;
%Fitting parameters
Vary1.g = [0.0 0.0 0.0]; %don't vary g
Vary1.lw = 0.;
Vary2.g = [0.0,0.00];
Vary2.lw = 0.;
Vary3.g = [0.00 0.00];
Vary3.lw = 0.00;
Vary1.weight = 0.05;
Vary2.weight = 0.05;
Vary3.weight = 0.05;

%calling fitting function

esfit('pepper',spc,{Sys1,Sys2,Sys3},{Vary1,Vary2,Vary3},Exp);
```

**S2B:** EasySpin script used to **simulate** the EPR spectrum of F229A $A_{red}$-$^{13}CO$, results shown in the main text, **Fig 5**. The simulation used the *pepper* function in Easyspin (total weight = 100%) to simulate individual components of the F229A $A_{red}$-$^{13}CO$.

```
clc; clear all;
%calls data
[x,y] = textread('1907152.txt'); %reads in experimental data for F229A
x = x/10; %x-offset
y=(y*11.25)+1; %arbitrary y-offset

%Component 1
Sys1.g = [2.0555 2.048 2.021]; %g-values O-conformer
Sys1.weight = 0.203;
Sys1.lw = 2.85024;

%Component 2
Sys2.g = [2.0169 2.065]; %g-values for A-conformer
Sys2.lw = 5.75413;
Sys2.weight = 0.597;

%Component 3  %g-valuesl for WT-Conformer
Sys3.g = [2.07825 2.031];
Sys3.lw = 7.775;
Sys3.weight = 0.20;

Exp.mwFreq = 34.85;      % GHz
Exp.Range = [1150 1248];% mT
Exp.Harmonic = 0; %absorption spectrum


Opt.Method = 'perturb';
Opt.nKnots = [61 0];
Exp.Harmonic = 0;

%Plotting

[xsim,ysim] = pepper({Sys1,Sys2,Sys3},Exp,Opt);
plot(x,y,xsim, ysim);

%Export

data = [xsim' ysim'];
save('ASCFull041120.txt','data','-ascii');
```

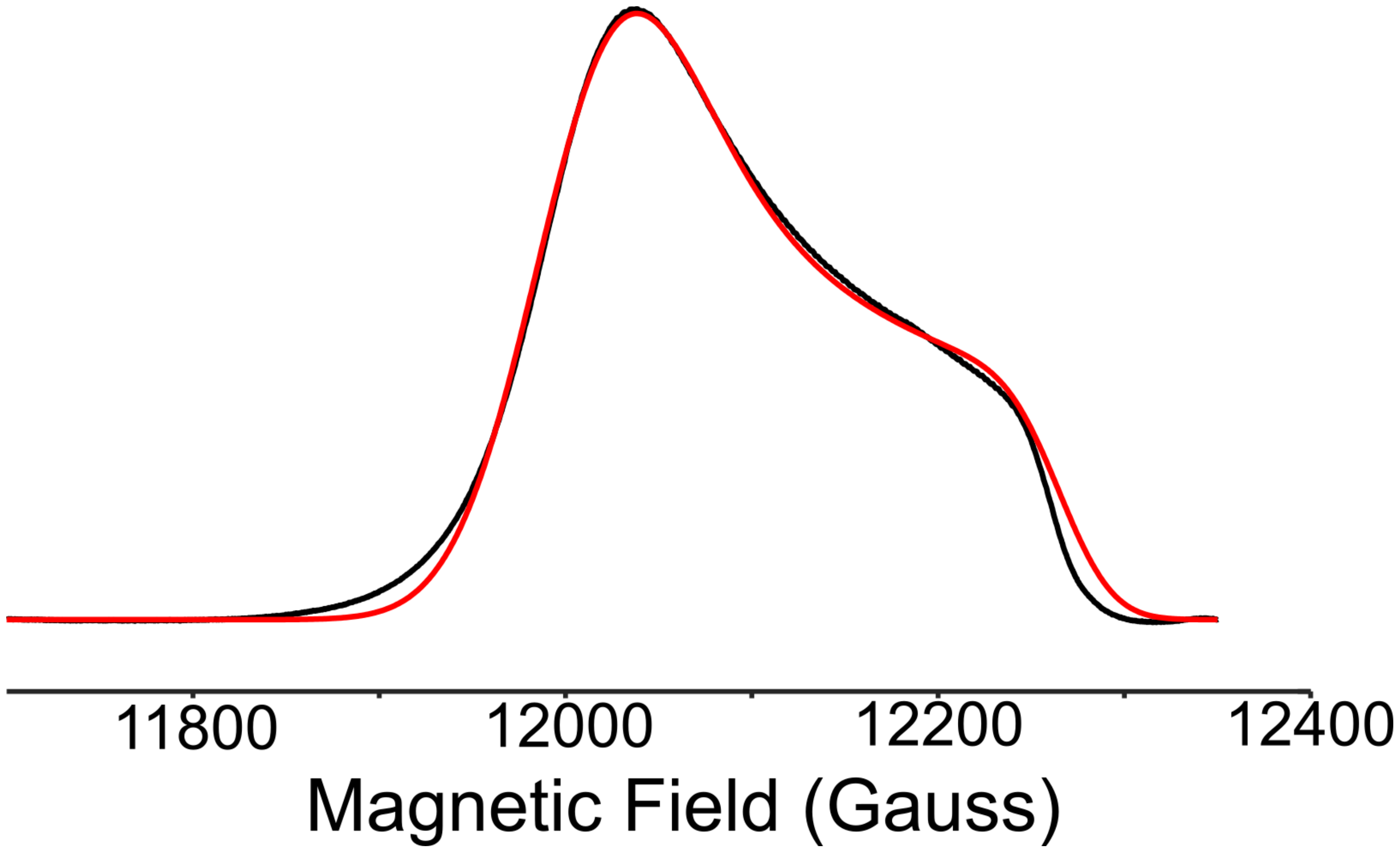


**Figure S3** simulation of the 35 GHz CW rapid-passage absorption-display EPR spectrum of F229W of $A_{red}$ -$^{13}$CO (Red). The simulation was prepared using the *pepper* function of the EasySpin simulation package. The g-values are listed in **Table S2.**

```
clc; clear all;

%calls data

[x,y] = textread('192310z1.txt','%f %f'); %reads in experimental data for F229W
x=x/10;
% Experimental Values
Sys.g = [2.0805 2.03675]; %F229W
Sys.lw = [5.25];

Sys.HStrain = [235.555 350 6.03435]

%Experimental Settings
Exp.mwFreq = 34.96;      % GHz
Exp.Range = [1180 1230];% mT
Exp.Harmonic = 0; %absorption spectrum
Exp.Temperature = 2; % 2K CW

Opt.Method = 'perturb';
Opt.nKnots = [61 0];
Exp.Harmonic = 0;

%plotting
[xsim,ysim] = pepper({Sys},Exp,Opt);

plot(x,y,xsim, ysim);
```

**Figure S4:** EasySpin script used to **simulate** the EPR spectrum of F229W $A_{red}$-$^{13}CO$. The simulation used the *pepper* function in Easyspin. The g-values are listed in **Table 1** in main text.

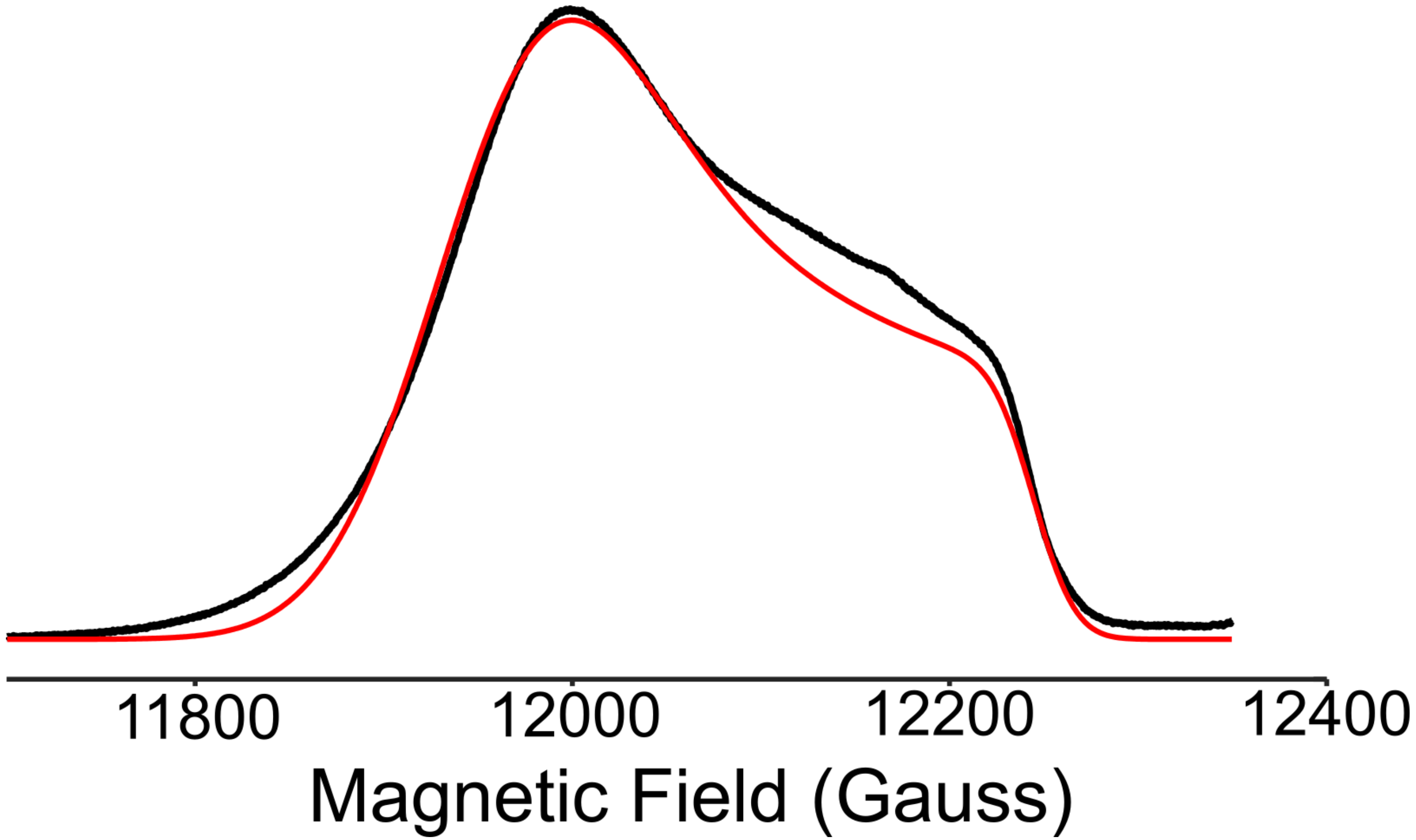


**Figure S5** simulation of the 35 GHz CW rapid-passage absorption-display EPR spectrum of WT F229 of $A_{red}$ -$^{13}$CO (Red). The simulation was prepared using the *pepper* function of the EasySpin simulation package and the spin-Hamiltonian parameters in **Table 1**. The intensity not accounted for by the simulation, at fields around ~12,150G, is associated with impurities; these features are more clearly seen in **Fig 1**, main text.

```
clc; clear all;

%calls data
[x,y] = textread('1906WT1.txt','%f %f'); %reads in experimental data for
F229W
x=x/10;
% Experimental Values
Sys.g = [2.085 2.03649]; %WT component
Sys.lwpp = [3.25];
Sys.A = [10 18];
Sys.Nucs = '13C';

Sys.HStrain = [354.908 82.0476 12.1184];

%Experimental Conditions
Exp.mwFreq = 34.89;      % GHz
Exp.Range = [1170 1235];% mT
Exp.Harmonic = 0; %absorption spectrum
Exp.Temperature = 2; % 2K CW

%Optimization
Opt.Method = 'perturb';
Opt.nKnots = [61 0];


%Plotting

[xsim,ysim] = pepper({Sys},Exp,Opt);

plot(x,y,xsim, ysim);
```

**Figure S6:** EasySpin script used to **simulate** the EPR spectrum of F229W $A_{red}$-$^{13}CO$. The simulation used the *pepper* function in Easyspin.
